# Real-time suppression and amplification of frequency-specific neural activity using stimulation evoked oscillations

**DOI:** 10.1101/2020.02.09.940643

**Authors:** David Escobar Sanabria, Luke A. Johnson, Ying Yu, Zachary Busby, Shane Nebeck, Jianyu Zhang, Noam Harel, Matthew D. Johnson, Gregory F. Molnar, Jerrold L. Vitek

## Abstract

**Background:** Approaches to predictably control neural oscillations are needed to understand their causal role in brain function in healthy or diseased states and to advance the development of neuromodulation therapies. In this study, we present a closed-loop neural control and optimization framework to actively suppress or amplify low-frequency neural oscillations observed in local field potentials in real-time by using electrical stimulation.

**Objective/Hypothesis:** The rationale behind this control approach and our working hypothesis is that neural oscillatory activity evoked by electrical pulses can suppress or amplify spontaneous oscillations via destructive or constructive interference when stimulation pulses are continuously delivered with appropriate amplitudes and at precise phases of these oscillations in a closed-loop scheme.

**Methods:** We tested our hypothesis in two nonhuman primates that exhibited a robust increase in low-frequency (8-30 Hz) oscillatory power in the subthalamic nucleus (STN) following administration of the neurotoxin 1-methyl-4-phenyl-1,2,3,6-tetrahydropyridine (MPTP). To test our neural control approach, we targeted 8-17 Hz oscillations and used electrode arrays and electrical stimulation waveforms similar to those used in humans chronically implanted with brain stimulation systems. Stimulation parameters that maximize the suppression or amplification of neural oscillations were predicted using mathematical models of the stimulation evoked oscillations.

**Results:** Our neural control and optimization approach was capable of actively and robustly suppressing or amplifying oscillations in the targeted frequency band (8-17 Hz) in real-time in the studied subjects.

**Conclusions:** The results from this study support our hypothesis and suggest that the proposed neural control framework allows one to characterize in controlled experiments the functional role of frequency-specific neural oscillations by using electrodes and stimulation waveforms currently being employed in humans.

## Introduction

Neuromodulation approaches that predictably control circuit-level neural dynamics in real-time will be of utility in neuroscience to deductively infer causal relationships between controlled changes in these dynamics and brain function. These control approaches could also help identify neural processes causally linked to the manifestation of brain conditions and inform the development of neuromodulation therapies. Neural oscillatory dynamics observed from local field potentials (LFP) are of particular interest to the development of brain stimulation therapies that use feedback signals to adjust stimulation, given the long-term stability of LFP recordings in cortical and subcortical brain structures in human subjects[1,2]. Evidence from experimental studies and computer simulations suggest that LFP oscillations at low-frequency (<100 Hz) are generated predominantly by synchronized synaptic inputs to neuronal ensembles near the recording site[3,4]. Controlling synchronized synaptic activity in a targeted neuronal ensemble can therefore help modulate information flowing into the target, and thereby influence the information flowing out of the target. Feedback (closed-loop) control systems offer the ability to control neural oscillatory activity by using LFP activity as a feedback signal and electrical stimulation for actuation[5].

Closed-loop stimulation systems being tested in human subjects deliver isochronal (fixed-frequency) electrical pulses on demand based upon neurophysiological signals extracted from LFP recordings. These on-demand systems have been used in patients with epilepsy[6] and Parkinson’s disease (PD)[7–10] to minimize the amount of stimulation energy delivered to the brain and reduce the likelihood of side effects that may occur with continuous stimulation as a result of current spread beyond the targeted region. Current on-demand systems based on isochronal stimulation are, however, not capable of actively suppressing or amplifying neural oscillatory activity in <u>specific</u> frequency bands in <u>real-time</u>.

Stimulation timed to the phase of neural oscillations has been proposed as a feedback strategy to alter the dynamics of neuronal populations [11,12]. A recent experimental study has shown that brief administration of electrical pulses phase-locked to oscillations in cortical networks can alter plasticity in nonhuman primates[13]. A study with PD patients undergoing DBS implantation surgery suggested that open-loop, isochronal, low-frequency stimulation delivered dorsal to the subthalamic nucleus (STN) could alter the amplitude of low-frequency oscillations recorded in the STN, whenever consecutive stimulation pulses landed at specific phase angles of these oscillations[14]. A recent study with nonhuman primates has also shown that brief stimulation bursts of intra-cortical microstimulation aligned with the peaks or troughs of neural oscillations in the motor cortex can result in changes in the amplitude of LFP oscillations near the stimulating electrodes, neuronal firing, and motor performance when delivered during a neuro-feedback task[15]. While phase-locked stimulation holds promise for modulating oscillatory activity within a neuronal ensemble, an approach to actively and predictably suppress or amplify neural oscillations in the circuit-level by using implantable electrodes remains to be developed. This approach together with strategies to optimize stimulation parameters based on neurophysiological data need to be developed if we are to bring active neural control technologies to clinical studies.

In this study, we demonstrate that active suppression or amplification of spontaneous neural oscillations can be achieved in real-time by using destructive (or constructive) interference created with damped oscillations evoked by electrical pulses. The rationale behind this approach is that spontaneous oscillations resulting from synaptic inputs to a targeted neuronal ensemble can be actively overwritten using stimulation-evoked oscillations. We employed a feedback control and optimization framework in which LFP measurements from chronically implantable electrodes are used for both sensing and delivering macro-stimulation. In this framework, active suppression or amplification of neural oscillations is achieved by continuously delivering stimulation pulses of adequate amplitude at precise phases of these oscillations in a closed-loop scheme. This strategy enables the use of stimulation waveforms with fixed amplitude similar to those used in humans chronically implanted with brain stimulation systems, making this technique feasible for human studies in the near future. We solved the problem of experimentally finding optimal stimulation parameters to amplify or suppress neural oscillations by using mathematical models of stimulation evoked potentials and computer simulations of the closed-loop neuromodulation system. This neural control framework, referred to as Closed-Loop Stimulation-Evoked Interference (CL-SEI), was tested in two parkinsonian nonhuman primates, which exhibited a robust increase in the power of oscillations in the 8-30 Hz frequency band following administration of the neurotoxin 1-methyl-4-phenyl-1,2,3,6-tetrahydropyridine (MPTP), similar to the oscillations observed in PD patients and thought to underlie the development of PD motor signs[16,17]. We were able to actively suppress or amplify STN oscillations in the studied animals by delivering stimulation pulses in the internal segment of the globus pallidus (GPi). Our data also demonstrates that the mechanism by which CL-SEI exerts its modulatory effect is constructive or destructive interference between low-frequency oscillations evoked by stimulation and the spontaneous oscillations in the site targeted for modulation.

## Material and Methods

### Instrumentation and subjects

All procedures were approved by the University of Minnesota Institutional Animal Care and Use Committee (IACUC) and complied with United States Public Health Service policy on the humane care and use of laboratory animals. Two adult rhesus macaques (female Macaca mulatta) were used in this study: subject J (17 years) and P (18 years). Each animal was implanted in both the STN and GPi with 8-contact scaled-down versions of human DBS leads (0.5 mm contact height, 0.5 mm inter-contact spacing, 0.625 mm diameter, NuMED, Inc.) as previously described[18]. The target locations for implantation of both STN and GPi leads were confirmed via microelectrode recordings of single unit activity and following established mapping approaches [19–22]. Briefly, regions in the STN in the naïve animals were characterized by tonic neuronal firing with 19-25 spikes/s, changes in firing rates associated with passive manipulation of the upper and lower limbs, and the contrast with its anterior and lateral borders. The GPi of the naïve animals was localized by identifying tonic discharges (10-107 spikes/s) with little pauses and frequent irregularities in the firing rate; defining the dorsal, posterior and lateral borders of GPi; and identifying the optic tract at the ventral border of GPi. We have previously described this approach in detail and refer the reader to Vitek et al (1998) [20]. DBS localization was confirmed post-mortem via histological reconstructions obtained by removing coronal (animal J) or sagittal (animal P) slices of 40-micron thickness and taking photographs of the brain after each slice was removed. The position of the DBS leads based to the histological reconstructions is shown in Fig. A.1. Following implantation and recordings in the naïve condition, the animals were rendered parkinsonian by systemic-intramuscular and intra-carotid injections of the neurotoxin MPTP [23]. Subject J received three intramuscular injections and one intra-carotid injection (0.3-0.4 mg/kg each, total 1.4 mg/kg). Subject P received weekly or biweekly systemic intramuscular MPTP injections to progressively induce mild, moderate, and severe parkinsonian states (total dose of 11.65 mg/kg).

### Experimental timeline

Neurophysiological data in the resting state of the studied subjects were collected several days before and after induction of parkinsonism. Data for animal J are from 20 sessions (19 days) in the normal state and 21 sessions (16 days) in the parkinsonian state. Data for animal P are from 5 sessions (5 days) in the normal state and 12 sessions (12 days) in the parkinsonian state. We used these datasets to characterize how neural oscillations across frequencies changed in the STN from the normal to the parkinsonian condition and to identify oscillations targeted for modulation using CL-SEI. Data to characterize evoked oscillations were collected in the parkinsonian condition of the subjects during the recording sessions dedicated to characterize the CL-SEI technique. Experiments to tune and validate artifact removal and signal processing algorithms were followed by experiments to characterize the effect of CL-SEI on neural oscillatory activity. Experiments with animal J provided us with neurophysiological data to validate the proposed closed-loop stimulation methodology. Due to complications with the animal’s medical condition it had to be euthanized and a second animal (P) was used to characterize optimal parameters of CL-SEI to amplify or suppress neural oscillations in a targeted frequency band. Experiments in animal P started only after the animal was in a severe parkinsonian state and we were unable to objectively quantify the behavioral effects of CL-SEI.

### Neural data acquisition

Resting state, off-stimulation **n**eural data were collected using a Tucker David Technologies (TDT, Alachua, FL, USA) workstation with a sampling rate of ~24 KHz. Signals were band-pass filtered (0.5-1400 Hz) to extract LFP data and down-sampled to ~3 KHz for analysis. LFPs were created by subtracting potentials from contacts estimated to be within the STN to suppress far field potentials. We verified that the animals were alert using video monitoring and image processing techniques[18]. The power of neural oscillations across frequencies was computed using power spectral densities (PSDs) and the Welch’s method using independent data segments of 15 seconds. A total of 185 data segments in the normal state and 312 in the parkinsonian state were analyzed for animal J. In animal P, 96 datasets in the normal state and 136 in the parkinsonian state were analyzed. The PSDs were computed by using 214 points in the Fast Fourier Transform, a Hamming window of 1.34 sec (¼ 214 points), zero padding (¾ 214 points), and an overlap of 50%. The resulting frequency resolution was 0.19 Hz. The PSD curves were normalized with respect to the total power, which is the sum of PSD values over all frequencies. Oscillatory modes with the highest power across frequency were selected for modulation using CL-SEI. Analyses were performed using customized scripts in MATLAB (Mathworks, Natick, MA, USA).

### Stimulation evoked potentials

Electrical stimulation was delivered in the GPi to modulate oscillations in the STN for two reasons. First, large evoked potentials were observed in the STN of both animals when stimulation (bipolar, biphasic) was delivered in the GPi. Second, stimulation in the GPi resulted in smaller stimulation-induced artifacts in the STN than those observed when stimulation was delivered within the STN itself, facilitating the suppression of artifacts in real-time. The external segment of the globus pallidus (GPe) was not considered as a target for stimulation in this study because subject P did not have more than one contact spanning this brain structure and the GPi is a more standard target for stimulation in clinical practice. We characterized the temporal dynamics of stimulation-evoked potentials in the STN by delivering low-frequency (<4 Hz) anodal and cathodal stimulation to the GPi, computing stimulation triggered averages of the LFP time series in the STN for both anodal and cathodal stimulation (*X*_*a*_(*t*) and *X*_*c*_(*t*)), and calculating the mean between the anodal and cathodal responses ((*X*_*a*_(*t*) + *X*_*c*_(*t*))/2). This mean evoked response attenuates the stimulation artifacts while preserving the neural evoked potential[24]. The evoked potentials were computed with at least 83 data segments in animal J and 482 data segments in animal P. Cathodal and anodal responses were aligned to the samples at which transition from negative to positive (or from positive to negative) phases of the stimulation pulses were detected. This alignment helps cancel out the stimulation-induced artifacts, but introduces a phase lag of ~80 us between the cathodal and anodal responses. This phase lag is negligible because of the much larger period of the evoked oscillations. See Fig. A.2. To eliminate residual artifacts, the time series were linearly interpolated in periods where the residual artifacts were observed. In animal J, the current source and sink of the stimulation system were connected to contacts C3 and C4 of the DBS lead, respectively. Contacts are labeled C0-C7 with C0 being the most ventral contact. In animal P, the current source and sink were connected to contacts C4 and C6 of the lead, respectively. The cathodal current source stimulation that we employed consists of delivering a symmetric biphasic pulse with a negative phase followed by a positive phase. In the anodal stimulation, a positive phase is followed by a negative phase.

The amplitude of the stimulation pulses used to characterize evoked potentials and implement the CL-SEI technique in both animal J and animal P was below 800 μA, which was below the stimulation threshold at which we observed muscle contractions associated with activating the internal capsule. These muscle contractions are the result of activating corticobulbar or corticospinal pathways that in turn activate cranial or spinal motor neurons [25]. The side effect threshold of each subject was based on assessments with high-frequency (130 Hz), isochronal, bipolar, cathodal stimulation performed as a standard procedure in our laboratory to establish the maximum amplitude that can be delivered to each subject. In these assessments, muscle contractions of the subject’s mouth and limbs were monitored at different current intensities at a fixed pulse-width and frequency. The smallest current amplitude that resulted in muscle contraction was set as the threshold for each subject. These thresholds were likely conservative for our study as the number of pulses per unit time delivered to the brain via CL-SEI is smaller than that associated with isochronal stimulation delivered at 130 Hz.

### Implementation of neural control approach

We implemented CL-SEI using a TDT neurophysiological recording and stimulation system (TDT, Alachua, FL, USA) and a computer fully dedicated to executing signal processing and control algorithms in real-time (Simulink Real-Time, Natick, MA, USA). LFP data were sampled at ~24 KHz in the neurophysiological recording system, amplified in the +/−10 V range, and transmitted to the control computer via analog channels. Stimulation was delivered using bipolar, charge-balanced, biphasic, symmetric stimulation pulses with pulse widths of 80 µs. The cathode and anode of the stimulation system were connected to separate contacts in the DBS lead located within the GPi as described earlier. The STN LFPs were digitized in the control computer at ~24 KHz to reconstruct the shape of electrical artifacts induced by stimulation and minimize the effect of antialiasing filters on the measured artifacts. Electrical artifacts were suppressed from the LFP data by holding for 2.3 ms the value of the LFP sample acquired before the stimulation pulse was triggered. The 2.3 ms holding period was longer than the duration of the artifact transient but short enough to avoid degrading information recorded at low-frequency. The artifact transients and holding time were visually inspected using both raw data and the average artifact computed as (*X*_*a*_(*t*) − *X*_*c*_(*t*))/2, where *X*_*a*_(*t*) and *X*_*c*_(*t*) are the average anodal and cathodal responses to a stimulation pulse. The artifact-suppressed LFP data were then band-pass filtered in the frequency band selected for modulation using second-order Butterworth filters and subsequently down-sampled at ~3 KHz. A frequency range of 6 Hz around the targeted center frequencies was selected for the implementation of CL-SEI to allow for variations in the frequency of oscillatory activity while attenuating information outside the frequency band of interest.

The instantaneous phase and amplitude of oscillations in the targeted frequency band were estimated using a Hilbert transformer filter[26]. The Hilbert filter introduced 16 samples of time delay (5.3 ms) to the closed-loop interconnection. A train of stimulation pulses, three for animal J and one for animal P, were delivered at specified phase angles. whenever the amplitude envelope of the oscillations was above a selected threshold. In animal J, this threshold was equal to the 25th percentile of the oscillations’ envelope computed in the off-stimulation state. In animal P, this threshold was set equal to the 15th percentile of the envelope to enhance the suppression capabilities of CL-SEI. Thresholds were calculated in each experimental session (each day) using 60s of resting-alert state data without movement artifacts. Stimulation was turned off when the envelope was below the determined threshold to avoid delivering stimulation based on measurements with small signal to noise ratio. A train of 3 pulses (165 Hz intra-burst rate) was used in animal J to enhance the modulatory effect of evoked oscillations via temporal summation, with the first pulse of the train aligned to the targeted phase estimate. In subject P, both pulse trains (3 pulses at 165 Hz) and a single pulse were employed to test CL-SEI. The rationale behind using a pulse train was to increase the amplitude of evoked neural activity in the frequency targeted for modulation via superposition of oscillations generated by multiple stimulation pulses. The number of pulses in the train influences the amplitude and phase lag of the evoked oscillations in the targeted frequency band; hence, the optimal phase angle to suppress or amplify spontaneous oscillations using a pulse train is different from that associated with a single pulse. The total delay of the closed-loop interconnection, associated with hardware communication, was less than 1 ms.

### Estimation of optimal stimulation parameters

Mathematical models of the stimulation evoked oscillations were developed to estimate the stimulation phase and amplitude that maximize the suppression or amplification of neural oscillations. Evoked potentials were characterized using linear, time-invariant differential equations and parameterized via system identification techniques[27]. A dead-zone operator was included in the models as evoked potentials were observed when the stimulation amplitude was greater than or equal to a non-zero value (lower bound). A more detailed description of the evoked potential characterization is presented in the **Appendix A**. By using these mathematical models, numerical simulations were created in which the closed-loop algorithms were evaluated. The numerical simulations incorporated the same algorithms, filters, and time delays present in the real-time control computer to predict optimal stimulation amplitude and phase angles that suppress or amplify oscillations. In these computer simulations, evoked potential models as well as computer-generated sinusoidal oscillations (synthetic) or previously recorded LFP data were added to obtain the modulated oscillations. We used sinusoids with an amplitude equal to the mean amplitude of experimentally recorded STN oscillations as a synthetic LFP signal in the computer simulation. Stimulation was delivered at various phase angles (−175, −170, 165, …, 180 deg) and stimulation amplitudes (300, 350.…,750, 800 *μ*A) in the computer simulations and a search was performed to calculate the optimal phases and amplitudes that resulted in maximum suppression or amplification of the oscillations in the targeted frequency band. The mean amplitude envelope of the modulated signal in the targeted frequency band was then used to quantify the degree of modulation achieved using CL-SEI in the simulation.

### Quantification of neural modulation

We assessed whether the amplitude of neural oscillations changed when CL-SEI was delivered by using scalar measurements of the oscillations’ amplitude envelope. The analytical amplitude of neural oscillations in the targeted frequency band was computed by filtering the raw data in this band, applying the Hilbert transform, and calculating the magnitude of the analytic signal obtained from the Hilbert transform. The scalar observations are equal to the average of the amplitude envelope over non-overlapping windows of one-second duration and separated by 0.5 seconds windows. This separation time is greater than the maximum time between effectively independent data, calculated across subjects and conditions (off stimulation, amplification, suppression) by using the autocorrelation function [28,29]. The separation windows minimize the effect of serial correlations between observations that may arise due to filtering and slow fluctuations in the power of neural oscillations. See the Appendix and Fig. A.4 for a more detailed description of how we computed the time between effectively independent data. Pairwise differences between scalar measurements of the oscillations’ amplitude in two different conditions were assessed via the Wilcoxon rank-sum test. The p-values resulting from this test were corrected for the number of comparisons via the Bonferroni method. We assumed that the difference between measurements in the two conditions was significant when p < 0.05. The comparisons made were stimulation off vs amplification and stimulation off vs suppression. Effect sizes were assessed using the Cohen signed test (‘U3’), a non-parametric two sample test [30].

## Results

### Spontaneous neural activity can be modulated using CL-SEI

The studied animals exhibited a robust increase in the power of oscillations in the STN in the 8-30 Hz band following MPTP treatment. See PSD plots in **Figs. 1b,c**. The frequency bands with highest power and targeted for modulation were centered at around 11 Hz in animal J and 14 Hz in animal P (**Figs. 1b,c**). Potentials in the STN evoked by stimulation in the GPi exhibited short-latency, high-frequency oscillations as well as low-frequency oscillations (**Figs. 1d-e)**. The low-frequency components of the evoked responses were used in the CL-SEI system to modulate the targeted spontaneous oscillations in both animals.

**Figure 1.**
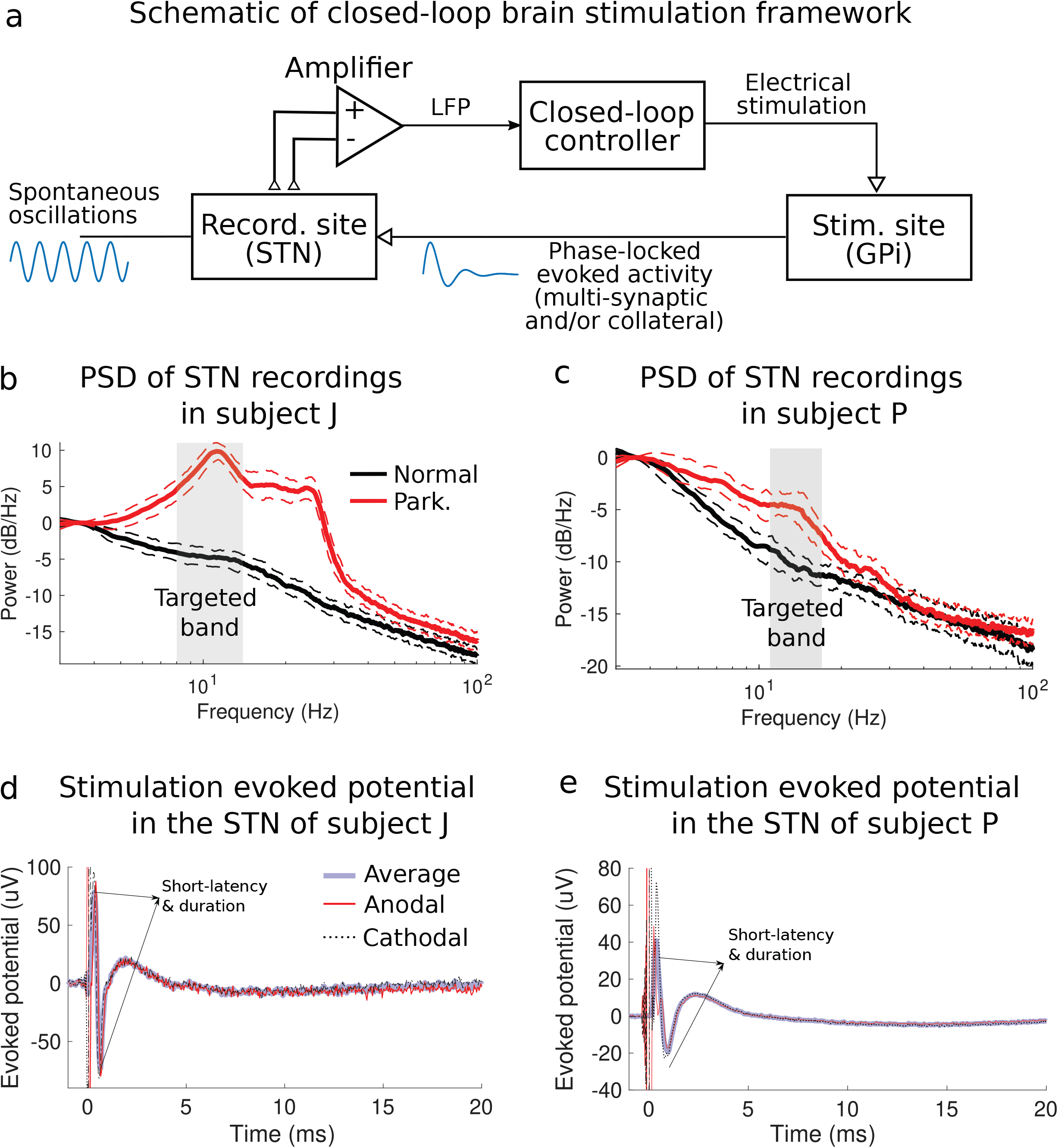
Closed-loop stimulation evoked interference (CL-SEI) framework and concept. (a) Schematic of closed-loop neuromodulation framework in which stimulation-evoked activity is used to suppress or amplify spontaneous neural oscillations when stimulations pulses of appropriate amplitude are phase-locked to these oscillations. The power spectral density (PSD) plots of spontaneous LFP recordings in the STN of the studied animal J and P in the normal and parkinsonian conditions are shown in (b) and (c), respectively. Shaded regions represent interquartile ranges of the PSD curves at each frequency. The frequency bands targeted for modulation using CL-SEI are highlighted in gray. These frequency bands have a range of 6 Hz with center frequencies equal to the peak frequency of the PSD plots. Electric potentials in the STN of animal J and P evoked by stimulation pulses in the GPi are shown in (d) and (e), respectively. The evoked potentials displayed in this figure are the average response of a single pulse with an amplitude equal to 600 μA for animal J and 800 *μ*A for animal P. Short-duration and short-latency evoked oscillations, depicted in (d) and (e), are associated with antidromic neural activation of the STN following the stimulus pulse.

By continuously delivering CL-SEI, we were able to modulate targeted neural oscillations in the STN of both animals. The amplitude of neural oscillations in the targeted frequency bands was a function of the phase angle at which stimulation pulses of fixed amplitude was delivered (**Figs. 2a-d**). The phase angles for which maximum amplification or suppression of neural oscillations were achieved at a fixed stimulation amplitude were unique, with a phase lag of 180 deg. The amplitude of oscillations in the modulated frequency band was significantly decreased (or increased) when stimulation was delivered at the optimal phase for suppression (or amplification) as compared to the amplitude when stimulation was turned off. In animal J, p=3.2e-9 and Cohen’s U3=0.98 for the amplification vs off-stimulation comparison, and p=1.7e-2 and U3=0.41 for the suppression vs off-stimulation comparison. In animal P, p=3.1e-48 and U3=0.97 for the amplification vs off-stimulation comparison, and p=1.2e-9 and U3=0.45 for the suppression vs off-stimulation comparison. Boxplots that summarize the effect of suppression and amplification of neural oscillations are shown in **Fig. 3**. These statistics indicate that the degree of amplification was higher than the degree of suppression in the experiments. One reason for this difference is that stimulation was discontinued when the amplitude envelope of the oscillations was below a prescribed threshold (Methods section).

**Figure 2.**
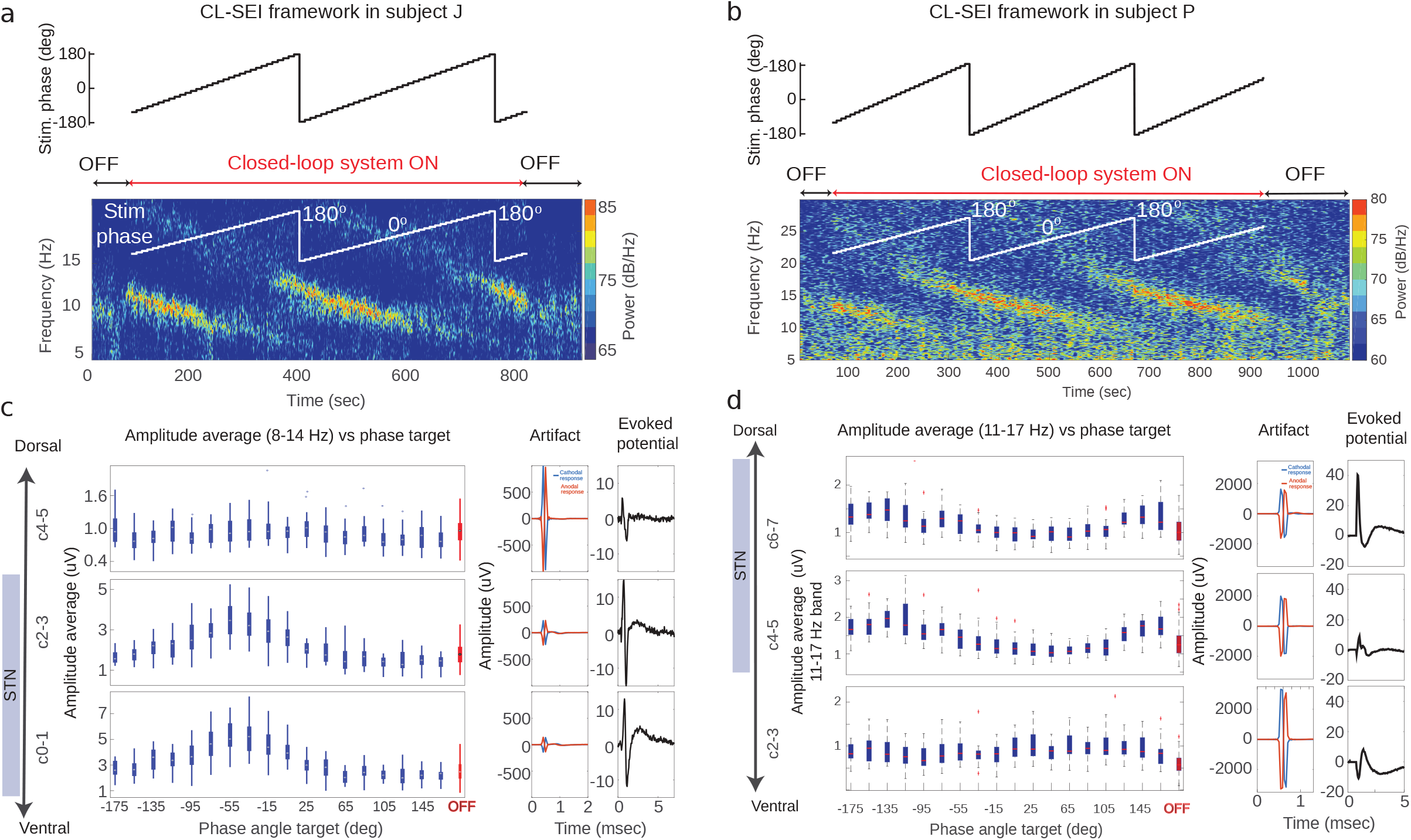
Amplitude of oscillations as a function of stimulation phase. (a, b) Results of experiments in which CL-SEI was tested in animal J and P. The targeted (reference) phase angle for stimulation was varied by 10 degrees every 10 seconds, while the stimulation current delivered to the GPi was fixed at 600 *μ*A in both animals. Spectrograms illustrating the power over time and frequencies for different phase angles in the OFF and ON stimulation state are shown in (a) and (b) for animal J and P, respectively. The spectrograms show data recorded using electrode pairs used to implement the closed-loop stimulation algorithm. The amplitude of neural oscillations as a function of the targeted phase angle and across electrode pairs is shown for animal J and P in (c) and (d), respectively. Electrodes C1-C3 in animal J and C4-C6 in animal P used for sensing are located within the STN. The artifacts and evoked potentials computed across electrode pairs are shown in (c) and (d) (center and right columns). Evoked potentials were calculated based on stimulation-triggered averages of LFPs by using both cathodal and anodal stimulation (*X*_*a*_(*t*) and *X*_*c*_(*t*)). These evoked potentials were calculated as ((*X*_*a*_(*t*) + *X*_*c*_(*t*))/2).

**Figure 3.**
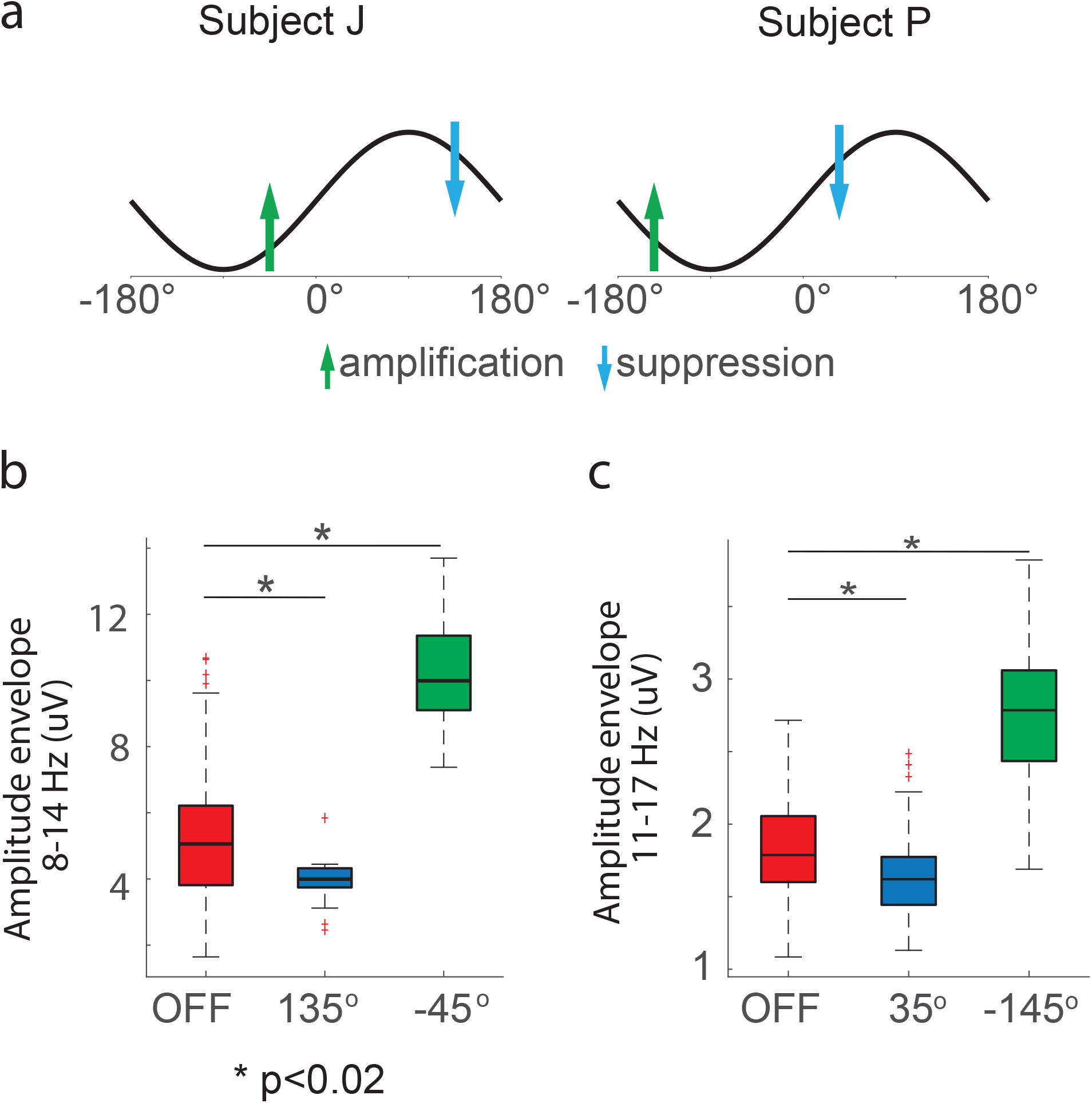
Differences between the amplitude of oscillations when stimulation was OFF and when stimulation was delivered at the phase angles associated with maximum suppression/amplification. (a) Schematic of phase angles at which stimulation was delivered to achieve maximum suppression (blue arrow) or amplification (green arrow) of neural oscillations in the 8-14 Hz band for animal J and 11-17 Hz band for animal P, respectively. (b) Box plots illustrating the interquartile ranges and medians of 1-second segments of the amplitude envelope of LFP data filtered in the targeted frequency band in the OFF stimulation state and when stimulation was delivered at phase angles that achieved maximum suppression and amplification of oscillations. We assumed that the difference between measurements in the two conditions was significant when p < 0.05. The p-values resulting from this test were corrected for the number of comparisons via the Bonferroni method. The number of data segments (*n*) the in the OFF stimulation, amplification, and suppression conditions for animal J were 143, 27, and 18, respectively; for animal P these numbers were 150, 20, and 20, respectively. The boxplots were created based on data from the experiment illustrated in **Fig. 2**.

### Modulation was local and not artifactual

In both subjects, the degree of modulation across different contact pairs in different regions inside and outside of the STN was not correlated with the size of the stimulation-induced artifacts, indicating that simulation-induced artifacts did not influence the observed modulation (**Figs. 2c,d**). Evoked potentials measured with electrodes estimated to be within the STN were larger than those outside the STN, suggesting that evoked neural activity was generated by neural sources and sinks located near electrodes located within the STN. Furthermore, spontaneous oscillations in the 8-30 Hz exhibited larger amplitudes inside the STN than outside (OFF state in **Fig. 2c,d**). The degree of amplification or suppression of these spontaneous oscillations achieved using CL-SEI was larger in electrodes located within the STN. Impedance measurements were consistent across sensing (STN) and stimulation (GPi) electrodes contacts in the DBS leads. A table is presented in the Appendix with the impedances of the STN and GPi lead contacts measured relative to a screw-electrode inserted in the skull of subject J and P. These impedances were measured immediately before the experiments presented in Figs. 2c-d began.

### Data-driven mathematical models enable estimation of optimal stimulation parameters and indicate that modulation was medicated by stimulation-evoked oscillations

The input-output relationship between the stimulation pulse and evoked potential was accurately characterized by the mathematical models described in the Methods Section and **Appendix A**(**Figs. 4a,b** and **Fig. A.2)**. A computer simulation of the experiment shown in Fig. 4d (same Fig. 2a) was created using LFP data in the stimulation-off condition and is shown in Fig. 4e. In both the experiment and simulation, a train of three pulses (165 Hz rate) was delivered. The computer-generated data reproduces the change in oscillatory power as a function of the stimulation phase observed in the in-vivo experiment (Fig. 4d vs Fig. 4e). This similarity indicates that 1) stimulation-evoked oscillations are responsible for the modulation in the CL-SEI framework, 2) linear superposition between evoked and spontaneous oscillations is a reasonable approximation for the interaction between these two components, and 3) delivering pulse trains results in temporal summation of evoked oscillations. To further illustrate how components of the evoked response in the targeted frequency band are involved in the modulation of spontaneous oscillations and how pulse trains enable temporal summation of this modulatory effect, we created computer simulations in which pulse trains and single pulses were delivered at 11 Hz in the absence of any spontaneous oscillatory activity (Fig. 4f). These simulations show that by using periodic pulse trains one can increase the amplitude of simulation-evoked oscillations in the targeted frequency band, which is the result of linear superposition of the evoked responses to individual pulses. The data shown in Fig. A3 confirm that temporal summation can be attained experimentally under these conditions. Note that due to the superposition principle described above and given that the periodic evoked response can be approximated as an infinite sum of sinusoidal components (i.e. Fourier transform), one can conclude that the evoked oscillatory components (Fig. 4f) in the modulated band are responsible for the modulation observed in both the computer simulations and experiments.

**Figure 4.**
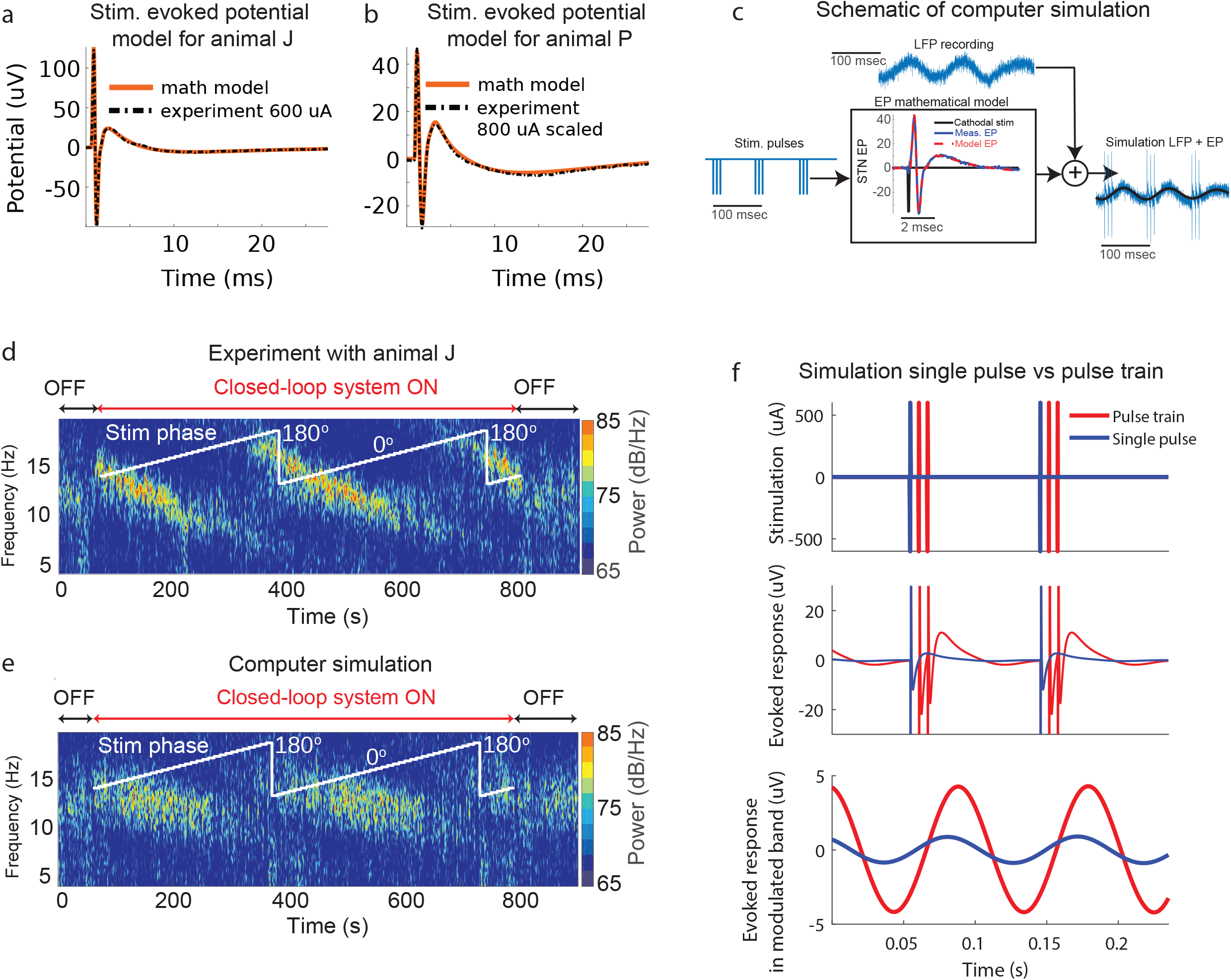
Computer simulations of CL-SEI. (a, b) Stimulation-evoked potentials calculated using experimental data together with their corresponding mathematical models. The evoked potential mathematical models are computed through the convolution of an evoked potential transfer function obtained via system identification techniques and the stimulation pulse mathematical function. (c) Schematic of computer simulation in which pre-recorded or computer-generated (synthetic) oscillations are added to the evoked potential (EP) time series to model the steady-state of modulated oscillations. The evoked potential time series is computed through the mathematical model with an input equal to stimulation pulses phase-locked to the oscillations in the selected frequency band. (d) Experimental data with animal J in which phase angles were varied over time. (e) Computer simulation of CL-SEI created using STN LFP data (stimulation-free) from animal J and mathematical models of the stimulation-evoked potentials. The targeted phase angles in both experiments and computer simulation were varied over time, increasing the phase angle by ten degrees each ten seconds. (f) Computer simulations of the evoked response to a single pulse and a train of three pulses delivered at the modulated frequency (11 Hz). This simulation was created using mathematical models of the stimulation evoked response of animal J. The top plot shows the single pulse and pulse train (165 Hz intra-burst rate) stimulation sequences used in the simulation. Both single pulses and stimulation trains were delivered periodically at 11 Hz. The middle plot shows the unfiltered, steady-state evoked response to both single pulses and pulse trains. The bottom plot shows the components of the evoked response filtered (forward and backward) in the 8-14 Hz band using a second order Butterworth filter. Evoked oscillatory components in the modulated band are larger for the stimulation trains than for single pulses.

We applied the search approach described in the Methods Section to estimate the optimal stimulation phase and amplitude to suppress/amplify 11-17 Hz oscillations in animal P by using one single stimulation pulse. The optimization considered the mean amplitude of the oscillations’ envelope measured in the resting, off-stimulation state of the animal. A map of the optimization search, with values of the mean amplitude of the oscillations’ envelope after CL-SEI is applied in the computer simulations at different phases and amplitudes, is shown in **Fig. 5a**. The optimal phase and amplitude to suppress oscillations were found to be 35 deg and 600 *μ*A, respectively. The optimal phase to amplify oscillations at the same amplitude (600*μA*) was found to be −145 deg. The optimal phase angles for both suppression (35 deg) and amplification (−145 deg) of oscillations at a fixed stimulation amplitude of 600 *μA* were the same as those found in the experiment depicted in **Figs. 2b** and **Fig. 3c.** This match suggests that the optimization approach can accurately predict the optimal stimulation parameters. Note that the optimal stimulation amplitude to <u>amplify oscillations</u> is equal to the maximum amplitude (upper bound) employed in the search since the size of the evoked oscillations is proportional to the stimulation amplitude[24,31]. The optimal amplitude to <u>suppress oscillations</u> is not necessarily equal to the optimal stimulation amplitude to amplify oscillations since evoked oscillations with high amplitude can create constructive interference even when delivering stimulation at phase angles at which suppression is achieved at lower stimulation amplitudes. The stimulation parameters found in the search algorithm were used in an experiment with the animal, in which we alternated between maximum suppression and amplification of neural oscillations (**Fig. 5b**). The results shown in **Fig. 5b** illustrate the capability of CL-SEI to actively suppress or amplify neural oscillations in real-time. Our experiments also show that while we attempt to modulate neural oscillations in the 11-17 Hz band, there are changes in power in adjacent frequency bands. In the suppression condition, the period of time in which the amplitude of the modulated oscillation was above threshold decreased as compared to the above-threshold period in the off-stimulation condition. The period in which the amplitude envelope of the oscillation was above threshold increased when the oscillations were amplified. The distributions of time periods in the off stimulation, suppression and amplification condition are shown in Fig. 5c.

**Figure 5.**
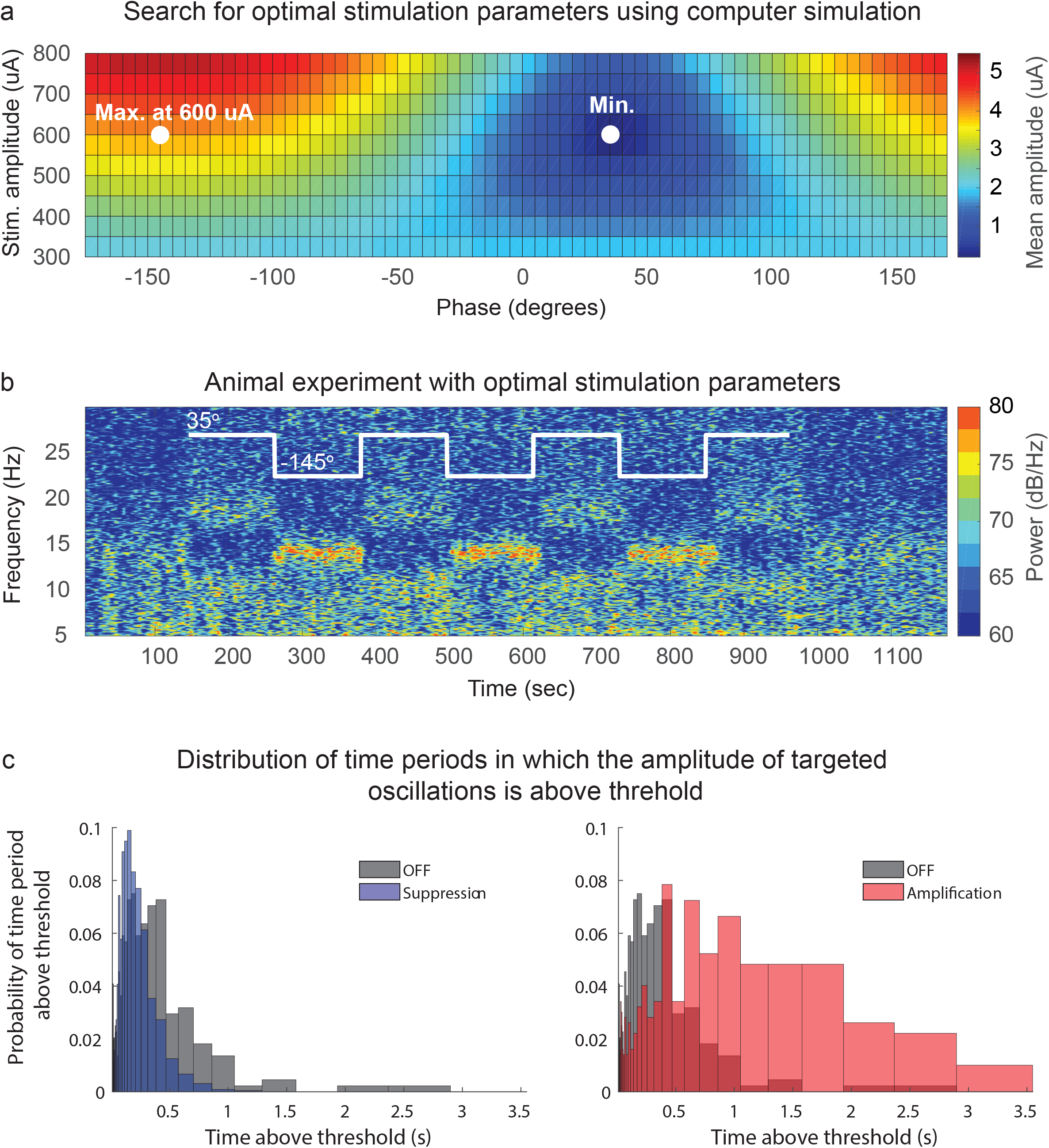
Optimization of CL-SEI parameters and experimental results. (a) Optimization map to estimate optimal stimulation amplitudes and phases to suppress/amplify neural oscillations in the 11-17 Hz band with a mean amplitude of 1.95 *μ*A in animal P by using CL-SEI. The color scale represents the mean amplitude of oscillations created in computer stimulations at different stimulation phases and amplitudes. The optimal phase and amplitude to suppress oscillations were found to be 35 deg and 600 *μ*A, respectively. (b) Spectrogram of STN LFP data in which CL-SEI was delivered at stimulation amplitude found to be optimal for suppression of 11-17 Hz oscillations (600 *μ*A). The stimulation phase was alternated between the optimal phase for suppression and the optimal phase for amplification at 600 *μ*A, which is illustrated with the white curve on the spectrogram. The recordings shown in this figure are from the electrode pair that was used for sensing in the closed-loop stimulation system. (c) Distribution of periods in which the amplitude of targeted oscillations was continuously above threshold in the off-stimulation, suppression and amplification conditions.

### Modulation was mediated by low-frequency evoked oscillations

Short duration (<3 ms) and short latency (<3 ms) components of the evoked potentials, likely associated with antidromic activation of the STN by stimulation of the GPi, were observed in our data (**Figs. 1d,e**), but have a negligible effect on the spectral measurements at low-frequency bands due to their low power in these bands. Our aforementioned observation that superposition between evoked and spontaneous oscillations explains the interaction between these two neural elements and implies that the frequency component of the stimulation-evoked response in the modulated frequency is what causes the modulatory effect on the spontaneous oscillations. The observed modulation (**Figs. 2a-d**) was therefore mediated by interference created by components of the stimulation evoked response in the modulated frequency band (low frequency). Our experimental and simulation data suggest that the same superposition principle enables us to enhance the amplitude of the evoked oscillations when pulse trains instead of pulses are delivered using the CL-SEI framework. See computer simulation in Fig. 4f and experimental data in Fig. A3.

## Discussion

We developed an experimental approach and optimization framework capable of precisely suppressing or amplifying the amplitude of neural oscillations in a specific frequency band with the resolution and time-scale required to characterize the functional role of oscillatory dynamics in brain circuits. The optimization framework described in this article resolves the time consuming problem of experimentally finding phase angles and stimulation amplitudes that maximize the degree of suppression or amplification of targeted neural oscillations. Finding the optimal stimulation phase and amplitude by conducting the experiment illustrated in Fig 2 for all stimulation amplitudes evaluated in the computer optimization (300, 350, …,800 uA) would take, in an ideal scenario and with a fully automated system, at least two hours (phase sweeps of 720 s duration for each stimulation amplitude). While feasible in animal experiments, this experimental search is intractable for studies involving human subjects. Together these neural control and optimization approaches enable researchers to address questions regarding the role of circuit-level, frequency-specific oscillatory dynamics in brain function in healthy or disease states. By improving our understanding of neural circuit dynamics in brain conditions, this closed-loop stimulation approach could also guide the development of subject specific neuromodulation therapies. The approach presented here can be applied to modulate oscillatory activity in either the healthy or disease state of a subject given that the following two conditions are satisfied: 1) the phase and amplitude of the targeted oscillations can be measured reliably, and 2) stimulation-evoked oscillations of sufficient power at the modulated frequency are observed in the same region where the spontaneous oscillations are measured. It is noteworthy that the role of oscillatory activity in disease states may be different from that in the healthy condition due to disease-related changes in brain circuitry. Additionally, the signal-to-noise ratio of the targeted oscillations may depend upon the state of the subject. For example, the large signal-to-noise ratio of low-frequency (8-30 Hz) oscillations observed in parkinsonian subjects facilitates the implementation of CL-SEI.

While we deliver stimulation based on the phase of neural oscillations, it should be noted that the rationale behind using phase feedback is different to that of methods that are intended to induce phase desynchronization across interconnected neurons[11,12], alter short-term plasticity[13], or modulate the balance between depolarization and hyperpolarization of neurons in the proximity of the stimulating electrodes[15]. The approach presented here differs in that it continuously overwrites the inputs of targeted neuronal populations to amplify or suppress oscillations via interference created with “new” oscillations evoked by electrical pulses. The main reason for employing phase-locking as a strategy to create neural interference is that it is more readily translated to clinical practice allowing one to use stimulation waveforms that do not change their amplitude, pulse-width, or shape in real-time. Phase-locked electrical stimulation with a fixed stimulation amplitude and pulse-width is, however, a suboptimal strategy to achieve interference because dynamic changes in the amplitude of neural oscillations are not considered in the optimization framework. Advanced feedback control algorithms that adapt to changes in the amplitude of oscillatory activity combined with stimulation systems that allow for instantaneous changes in stimulation parameters are more likely to produce better interference patterns and need to be developed if we are to achieve optimal control of pathological oscillatory activity in PD and other disorders via CL-SEI.

### Modulation mechanisms of CL-SEI

Our analysis indicates that modulation of spontaneous oscillations in the targeted frequency band stems from the superposition of these oscillations and components of the stimulation-evoked response in the targeted band. The mechanism by which CL-SEI exerts its effect on LFP oscillatory activity is likely the result of synaptic summation of stimulation-evoked synaptic inputs generated by activation of mono- or multi-synaptic connections to targeted neuronal population and spontaneous synaptic inputs to the same population [32–35]. We targeted oscillations in the 8-17 Hz band in the STN of two parkinsonian monkeys that exhibited an increase in power following MPTP administration. The mechanisms by which the amplitude of these oscillations increase in parkinsonian animals and are observed in PD patients is still being debated. Studies with the 6-OHDA mouse model of PD and computer simulations of the human STN indicate that a combination of synchronized inhibitory and excitatory synaptic inputs from the cerebral cortex and the external segment of the globus pallidus (GPe), innervating common neuronal ensembles contributes to the development of these oscillations in the STN[36,37]. In the experiments carried out in this study, we modulated neural oscillations in the STN in the 8-17 Hz band by delivering stimulation with appropriate amplitude in the GPi that was phase-locked to oscillations in the STN. The resultant modulation of oscillatory activity in the STN was mediated by low-frequency, stimulation-evoked oscillations during stimulation in the GPi. Although low-frequency damped oscillations in the STN evoked by GPi stimulation have not been characterized previously, they are likely associated with synaptic inputs to the STN[32–35]. Based on known anatomical connections, these evoked oscillations may emerge from a combination of one or more of the following: 1) antidromic activation of GPe followed by activation of collateral branches projecting to the STN[38]; 2) orthodromic activation of GPi-PPN-STN pathways, and 3) orthodromic activation of pallido-thalamo-cortico-subthalamic loops. Consequently, controlled changes in the amplitude of STN oscillations achieved via stimulation of the GPi may be mediated by simultaneous synaptic inputs to a number of different neuronal ensembles converging on the STN.

### Limitations

We assumed that spontaneous and evoked LFP oscillations measured in the STN were associated with synaptic inputs innervating common STN neuronal ensembles. This is a reasonable assumption given that STN neurons simultaneously receive inputs from multiple structures, including the GPe and motor cortex. However, we acknowledge that spontaneous and evoked oscillations may be generated by inputs to adjacent but different populations of neurons, and modulations observed in the experiments may be caused by the superposition of currents from different neuronal ensembles. Recordings of synaptic activity (postsynaptic potentials) and/or neuronal activity in populations of STN neurons during periods of CL-SEI are needed to characterize how this stimulation approach exerts its effect at the cellular level. The experimental data shown in **Figs. 2a,b** and **Fig. 5b** indicate that CL-SEI alters oscillatory activity in frequency bands not targeted for modulation. These alterations are, at least in part, due to the power content of the evoked neural activity in frequencies outside the band selected for modulation. Stimulation patterns different from fixed-amplitude pulse trains may be needed to reduce the activation of neural oscillatory activity in frequency bands outside that selected for modulation using CL-SEI. The experimental setup used in this study, with electrodes in both the STN and GPi, has been implemented in humans[39]; however, it is not standard practice as DBS leads are typically implanted in one brain target within the same cerebral hemisphere. In applications in which LFPs are recorded from and stimulation is delivered to the same brain region using a single electrode array (e.g. DBS for PD), hardware and/or software technologies that enable robust artifact removal are needed in order to rigorously implement CL-SEI. The relative benefit of CL-SEI compared to standard brain stimulation approaches using isochronal stimulation remains to be determined, but the potential utility of this technique for both improving current stimulation therapies and for understanding the pathophysiological basis underlying circuit disorders is compelling.

## CRediT Authorship Contribution Statement

**David Escobar Sanabria:** Conceptualization, Methodology, Software, Formal analysis, Validation, Investigation, Visualization, Writing – Original Draft, Supervision, Funding Acquisition. **Luke A. Johnson:** Resources, Investigation, Visualization, Writing – Review & Editing, Funding Acquisition. **Ying Yu:** Investigation (data collection). **Zachary Busby:** Investigation (data collection). **Shane Nebeck:** Investigation (data collection). **Jianyu Zhang:** Resources, Investigation. **Noam Harel:** Investigation (MR-CT image acquisition). **Matthew D. Johnson:** Writing – Review & Editing, Funding Acquisition. **Gregory F. Molnar:** Project administration, Supervision, Funding Acquisition. **Jerrold L. Vitek:** Conceptualization, Resources, Project administration, Supervision, Validation, Writing – Review & Editing, Funding Acquisition.

## Acknowledgments

We thank Tay Netoff for the discussions regarding closed-loop brain stimulation techniques as well as Claudia Hendrix, Devyn Bauer and Ben Teplitsky for their help with histology processing.

---

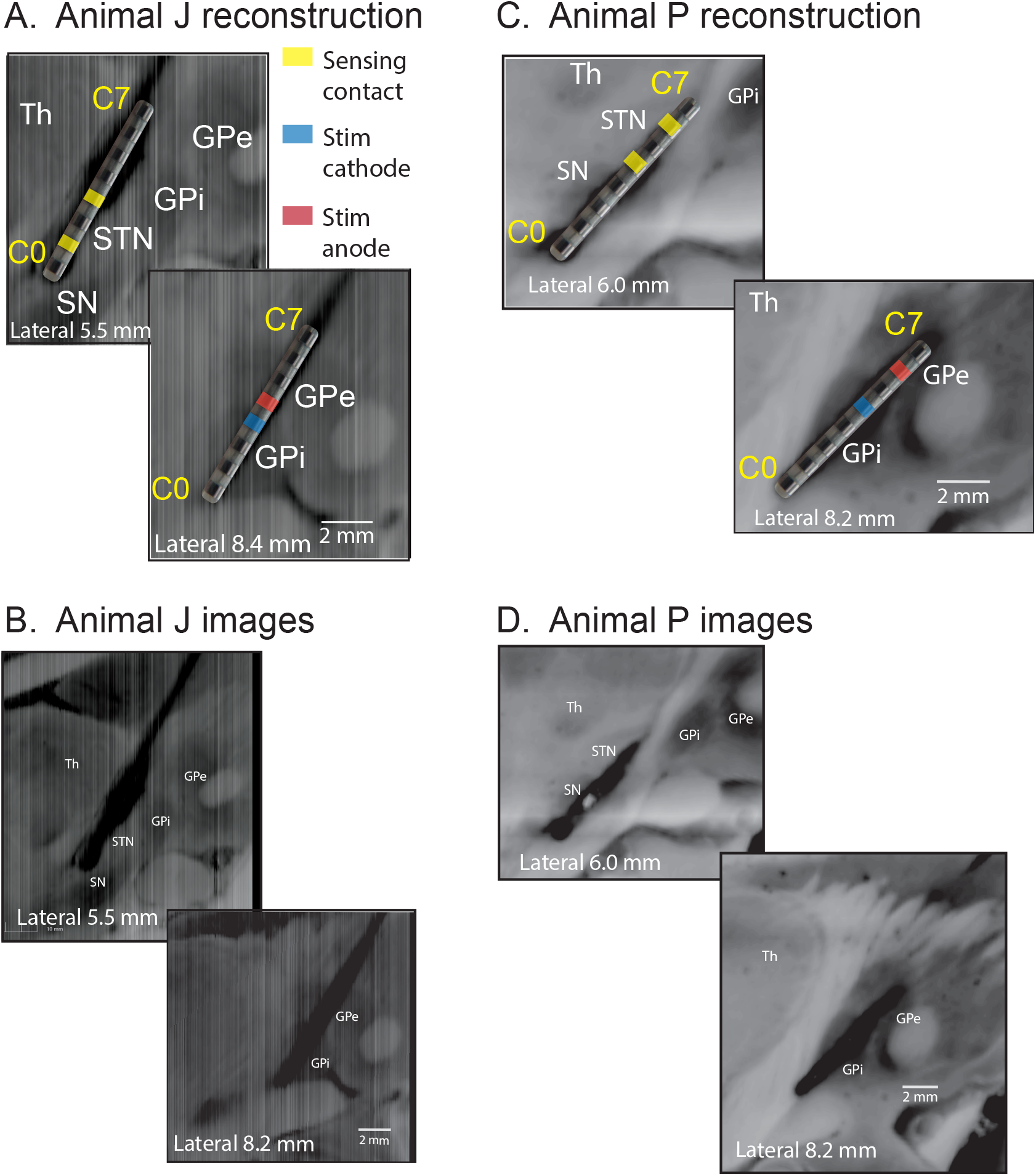

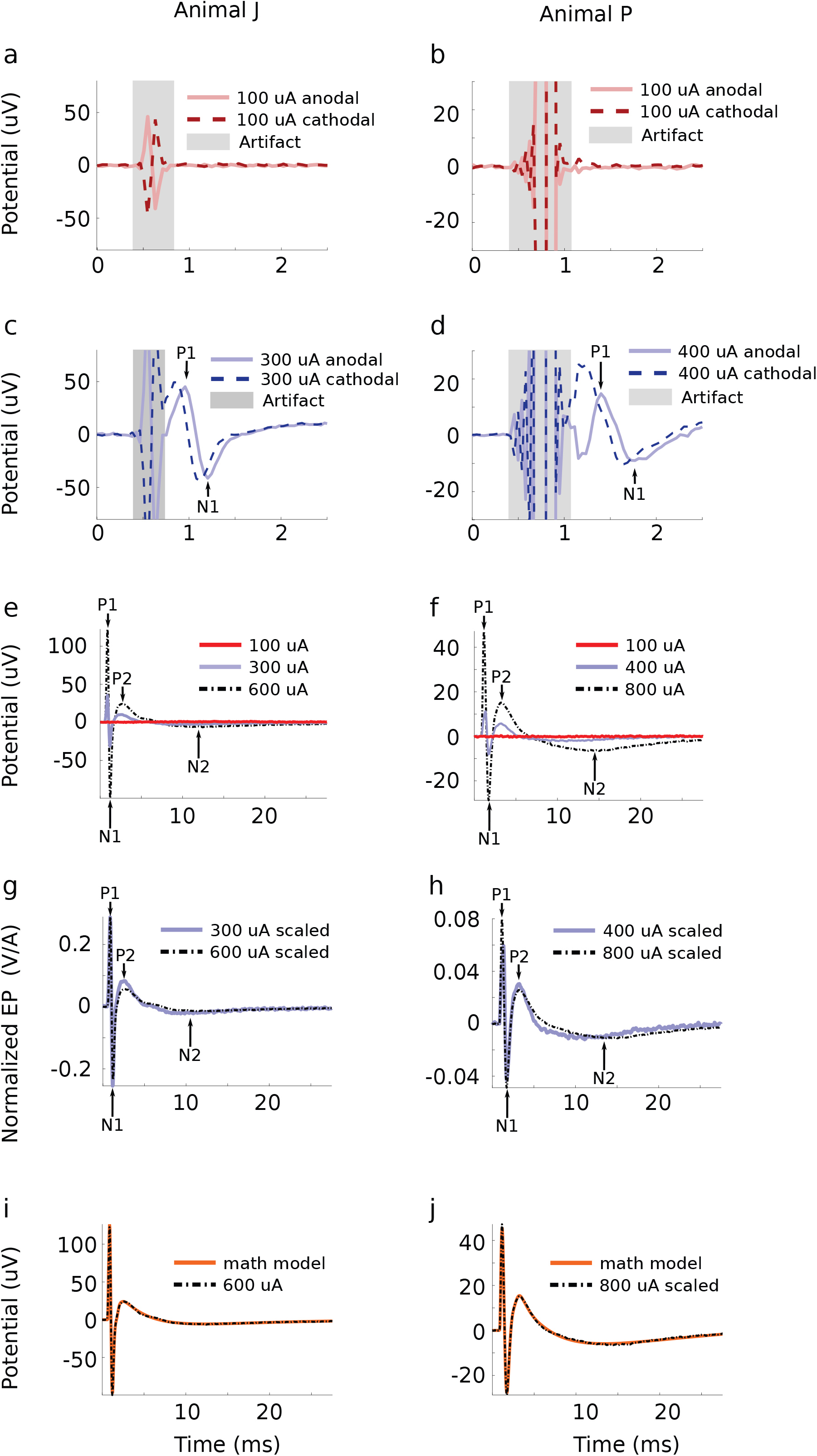

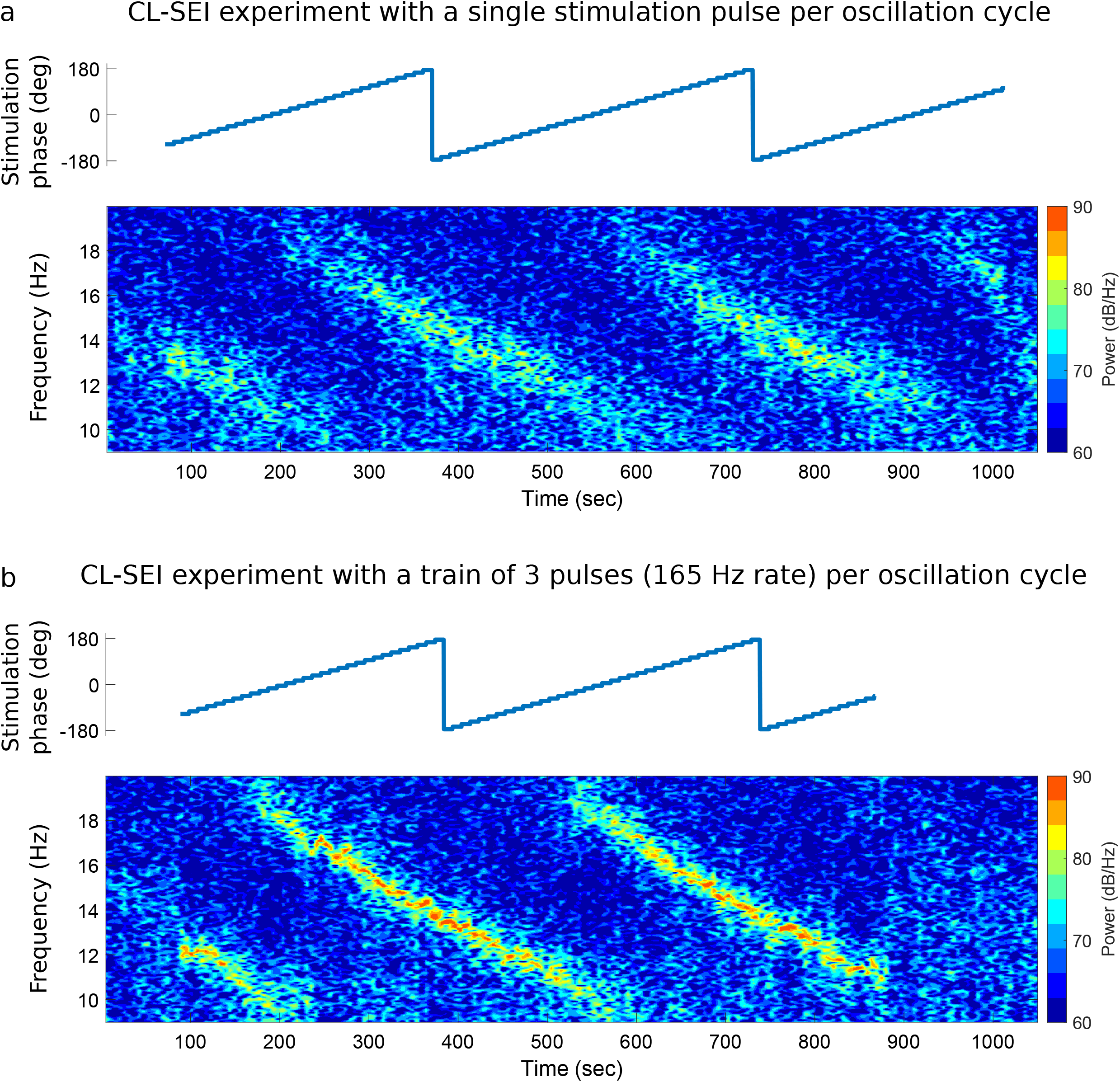

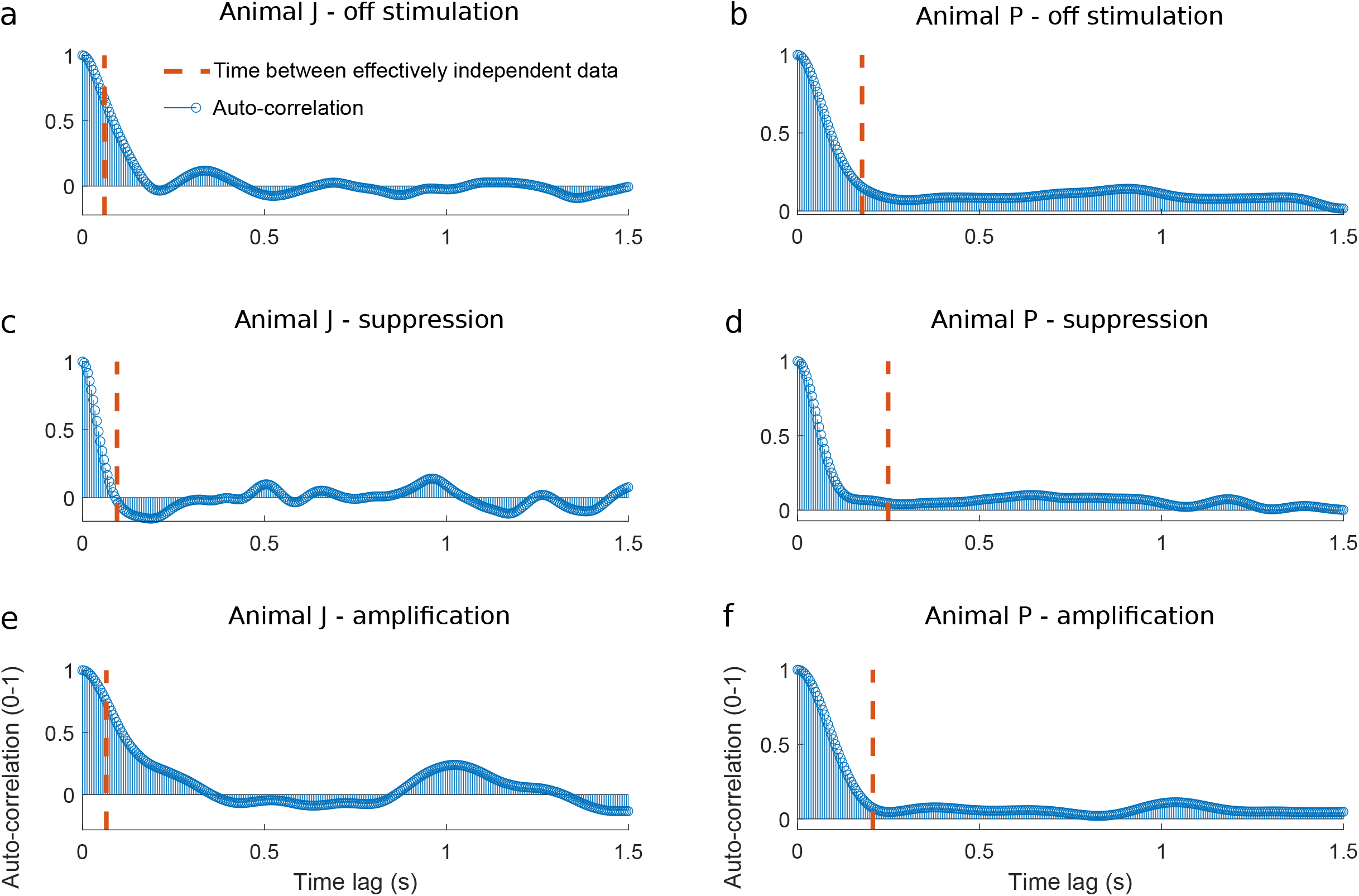

