## Appendix A for "Real-time suppression and amplification of frequency-specific neural activity using stimulation evoked oscillations"

#### Electrode Localization

The locations of the DBS electrode arrays implanted in the subthalamic nucleus (STN) and internal segment of the globus pallidus (GPi) were estimated using histological reconstructions in animal J and P (Fig. A.1). In animal J, electrodes C0 through C2 are within the STN and electrodes C2 through C4 are within the GPi. In this animal, the STN contacts selected for sensing were C1-C3 and the GPi contacts selected for stimulation were C3-C4. In animal P, electrodes C4 through C6 are within the STN and electrodes C3 through C6 are within the GPi. In this animal, the STN contacts selected for sensing were C4-C6 and the GPi contacts selected for stimulation were C4-C6

#### Characterization of Stimulation Evoked Oscillations

The amplitude of potentials in the STN evoked by stimulation in the GPi was characterized at three different stimulation amplitudes, in both anodal and cathodal configurations, in each animal (J and P). In both animals, stimulation evoked potentials were not observed for stimulation currents of 100 μA, either in the cathodal or anodal configuration. For stimulation currents of 300 μA and 600 μA in animal J, and 400 μA and 800 μA in animal P, evoked potentials were observed in both anodal and cathodal configurations (**Figs. A2a-f**). The absence of evoked neural activity for stimulation currents at 100 μA indicates that there is a minimum current for which evoked responses are generated by the electrical stimulation pulses. The evoked responses generated by anodal stimulation had a phase lag relative to those generated by cathodal stimulation, suggesting that the cathodal phase of the stimulus was responsible for the evoked response (**Figs. A2c,d**). The evoked potentials have a short-latency (<2 ms) and a short-duration (<2 ms) oscillation with peak P1 and trough N1 followed by oscillations of longer duration, including the oscillations with peak P2 and trough N2 (**Figs. A2c-f**).

The input-output relationship between the stimulation pulses and evoked potentials at sample time $k$ were characterized by the following mathematical model

$$y(k) = \begin{cases} [A-\bar{A}]\,(h_e * v)(k) & if\ A > \bar{A} \\ 0 & otherwise, \end{cases} \qquad \text{(Eq. A.1)}$$

where $h_e$ is a linear time-invariant transfer function estimated using instrumental variable system identification[1], $v(k)$ is the stimulation input described by the cathodal cycle of the stimulation pulse, the symbol $*$ is the convolution operation, $A$ is the stimulation current amplitude, $\bar{A}$ is the minimum stimulation current for which neural oscillations are evoked. The $A-\bar{A}$ factor is the effective stimulation gain for the linear system model when $A > \bar{A}$. The stimulation waveform is described by the equation

$v(k) = u(t) - u(t - T_{pw})$

$u(k) = \begin{cases} -1 & if\ k > 0 \\ 0 & otherwise, \end{cases}$

where $T_{pw}$ is the pulse width of the stimulation pulse and $u(k)$ the unitary step function. The constant scalar $\bar{A}$ is estimated using experimental data in which evoked potentials are calculated for two different stimulation current amplitudes in a range in which the amplitude of the evoked potentials scales linearly with the stimulation amplitude. Following the model structure of **Eq. A.1**, the evoked potentials generated by stimulation with two different stimulation amplitudes $A_1$ and $A_2$ have the following relationship at each sample $k$

$$\frac{\hat{y}(k, A_1)}{A_1 - \bar{A}} = \frac{\hat{y}(k, A_2)}{A_2 - \bar{A}}, \qquad (Eq. A.2)$$

whenever $A_1 > \bar{A}$ and $A_2 > \bar{A}$. We used a least squares minimization to estimate $\bar{A}$ based on **Eq. A.2** and using evoked potential time series $\hat{y}(k, A_1)$ and $\hat{y}(k, A_2)$ measured experimentally. For doing this estimation, we rearrange **Eq. A.2** as follows

$$Y\,\bar{A} = c, \qquad (Eq. A.3)$$

where $Y$ is a column vector of the form

$$Y = [\hat{y}(1, A_2) - \hat{y}(1, A_1), \hat{y}(2, A_2) - \hat{y}(2, A_1), \dots, \hat{y}(N, A_2) - \hat{y}(N, A_1)]^T,$$

with $N$ being the number of samples of the evoked potential time series. $c$ is a column vector of the form

$$c = [A_1\hat{y}(1, A_2) - A_2\hat{y}(1, A_1), A_1\hat{y}(2, A_2) - A_2\hat{y}(2, A_1), \dots, A_1\hat{y}(N, A_2) - A_2\hat{y}(N, A_1)]^T.$$

$\bar{A}$ is abstained via the least squares solution to **Eq. A.3**, which is given by $\bar{A} \approx (F^T F)^{-1} F^T b$.

The values of $\bar{A}$ for animal J and P are equal to 176 µA and 281 µA, respectively. The scaling of evoked potentials by a factor $A - \bar{A}$ is demonstrated in **Fig. A.2g-h**. The mathematical model of the evoked potential closely reproduces the evoked potential measurements in both animal J and P (**Fig. A.2 i,j**).

The discrete-time transfer functions $h_e$ are described by

$H_e(z) = \frac{b^T Z_b}{a^T Z_a} = \frac{[b_m, b_{m-1}, \dots b_1][z^m, z^{m-1}, \dots, z]^T}{[a_n, a_{n-1}, \dots a_1][z^n, z^{n-1}, \dots, z]^T}$, where $b$ is a column vector with real parameters and $Z_b$ a column vector with integer powers of $z$, the complex frequency representation of a discrete-time, linear, time-invariant system. The vectors $a$ and $b$ obtained via system identification for animal J are

$$a^T = [1.0,\ -8.160,\ 29.665,\quad -63.091, 86.563,\quad -79.499,\ 48.897, -19.431, 4.528,\ -0.472],$$

$$b^T = 1e-5 \cdot [0,\ 0.005,\ -0.038,\ 0.123\ -0.231\ 0.281\ -0.229\ 0.121\ -0.038\ 0.005],$$

and those for animal P are

$$a^T = [1.0, -7.893, 27.489, -55.355, 70.88, -59.674,\ 32.89, -11.367, 2.211,\ -0.181]$$

$$b^T = 1e-06\ \cdot [\,0.0005, -0.008, 0.06, -0.216, 0.416, -0.452, 0.2636, -0.061,\ -0.009, 0.0053].$$

### Optimization Approach

We calculated optimal stimulation amplitudes and phases for phase-locked stimulation to amplify or suppress neural oscillations by employing a search approach and the evoked potential mathematical models described in **Eq. A.1**. We modeled the spontaneous oscillations and phase-locked evoked potentials as

$$x_{raw}(k, A, \phi) = L\cos(\omega\, k\, T_s) + \sum_{m=1}^{M}\sum_{n=0}^{N} y(k - \left\lceil \frac{m}{\omega T_s} \right\rceil - \left\lceil \frac{\phi}{T_s} \right\rceil - n\, T_b, A)$$

where $\omega$ is the center frequency of the band targeted for modulation, $L$ is the mean amplitude of neural oscillations targeted for modulation and measured in the resting state of the studied subjects. $y$ is the function that characterizes the evoked potential given the stimulation amplitude $A$ and phase $\phi$. The term $T_s$ is the sampling period used in the discretization of time and $k = 1,2,\dots,K,$ is the sampled time. The term $T_b$ is the intra-burst period of a stimulation pulse train, and $N$ is the number of pulses in each pulse train. In this model, we set $M = \lfloor K\omega T_s \rfloor$ pulse trains, considering stimulation for all cycles of the cosine function $L\cos(\omega\, k\, T_s)$ in the interval from $k = 1$ to $k = KT_s$. The operations $\lfloor\cdot\rceil$ and $\lfloor\cdot\rfloor$ are the round and floor functions that transform a real number into an integer.

The component of $x_{raw}(k)$ in the targeted frequency band is given by the equation $x_{fil}(k) = (x_{raw} * h_{bp})(k)$, where $*$ is the convolution operation and $h_{bp}$ is the transfer function of a band-pass filter with cutoff frequencies delineating the targeted frequency band. We used second order Butterworth filters and bandwidth equal to 6 Hz. The focus of our optimization approach is to minimize (or maximize) the amplitude envelope of $x_{fil}$ by selecting a stimulation amplitude $A_{opt}$ and stimulation phase $\phi_{opt}$. This optimization is described by the expression

$$\min/\max_{\phi_m, A_q} \frac{1}{K}\sum_{k=1}^{K} Amp(x_{fil}(k)), \qquad (Eq. A.4)$$

where $Amp\big(x(k)\big) = |x(k) + jHT\big(x(k)\big)|$, $j = \sqrt{-1}$, HT is the Hilbert transform operator, $|\cdot|$ is the magnitude operation applied to a complex number. The phase angles $\phi_m$ are discretized between 0 and $2\pi$ so that $\phi_m = \frac{2\pi p}{\bar{P}}$, for $p = 0, 1, \dots, \bar{P} - 1$. The number of phase angles considered in our discretization was $\bar{P} = 72$, which leads to an angle resolution of 5 deg. The amplitude of the stimulation current was discretized by $A_q = A_0 + \frac{q(A_Q - A_0)}{Q}$, for $q = 0, 1, \dots, Q$. $A_0$ and $A_q$ are the lowest and highest values of stimulation amplitude used in the optimization. We used $A_0$=300 μA and $A_Q$ =800 μA for the optimization carried out for animal P as these currents were above $\bar{A}$ and below the threshold at which side effects associated with activating the internal capsule were observed. Q was set equal to 11, which gives us a stimulation amplitude resolution of 50 μA. The optimal values for the maximization or minimization (**Eq. A.4**) were obtained by doing a search over all discretized values of stimulation phase $\phi_p$ and amplitude $A_q$.

### Time Between Effectively Independent Data

The number of samples between effectively independent data, as described in [2,3], was computed as

$$N_{eq} = \frac{N}{1 + 2\sum_{\tau=1}^{N-1} \frac{N-\tau}{N}\,\rho(\tau)},$$

where N is the number of samples used in the calculation and $\rho(\tau)$ is the autocorrelation function estimated from the data. The time between effectively independent data is equal to $N_{eq}\Delta t$, where $\Delta t$ is the sampling period of the time series data. The time between effectively independent data for the amplitude envelope of neural oscillations in the off stimulation, suppression, and amplification conditions is presented in **Fig. A4** together with the autocorrelation functions. The time series used to calculate $N_{eq}$ and the autocorrelation function estimates shown in **Fig. A4** consisted of N=3,500 samples (10 sec).

#### Electrode Impedances

**Impedances at 1 KHz for sensing and stimulation electrodes (KOhms)**

| Subject J | | | | | | | | |
|---|---|---|---|---|---|---|---|---|
| **Contact** | **C0** | **C1** | **C2** | **C3** | **C4** | **C5** | **C6** | **C7** |
| **STN (sensing)** | 13.7 | 13.9 | 14.1 | 14.6 | 14.1 | 14.1 | 23.2 | 12.8 |
| **GPi (stimulation)** | 16.2 | 17.8 | 16.2 | 16.3 | 16.9 | 18.2 | 17.4 | 33.2 |

| Subject P | | | | | | | | |
|---|---|---|---|---|---|---|---|---|
| **Contact** | **C0** | **C1** | **C2** | **C3** | **C4** | **C5** | **C6** | **C7** |
| **STN (sensing)** | 7.4 | 7.2 | 6.8 | 6.3 | 6.3 | 6.4 | 6.6 | 6.7 |
| **GPi (stimulation)** | 6.3 | 6.2 | 15.6 | 7.4 | 6.3 | 6.7 | 6 | 7.9 |


**Impedances at 1 KHz for sensing and stimulation electrodes (KOhms)**

| Subject J | | | | | | | | |
|---|---|---|---|---|---|---|---|---|
| **Contact** | **C0** | **C1** | **C2** | **C3** | **C4** | **C5** | **C6** | **C7** |
| **STN (sensing)** | 13.7 | 13.9 | 14.1 | 14.6 | 14.1 | 14.1 | 23.2 | 12.8 |
| **GPi (stimulation)** | 16.2 | 17.8 | 16.2 | 16.3 | 16.9 | 18.2 | 17.4 | 33.2 |

| Subject P | | | | | | | | |
|---|---|---|---|---|---|---|---|---|
| **Contact** | **C0** | **C1** | **C2** | **C3** | **C4** | **C5** | **C6** | **C7** |
| **STN (sensing)** | 7.4 | 7.2 | 6.8 | 6.3 | 6.3 | 6.4 | 6.6 | 6.7 |
| **GPi (stimulation)** | 6.3 | 6.2 | 15.6 | 7.4 | 6.3 | 6.7 | 6 | 7.9 |


**Table A1**. Impedance measurement of DBS electrode arrays for subject J and P. The impedances were measured between each contact of the DBS lead and a screw-electrode inserted in the skull of the subjects. The impedances were measured at a frequency of 1 KHz.

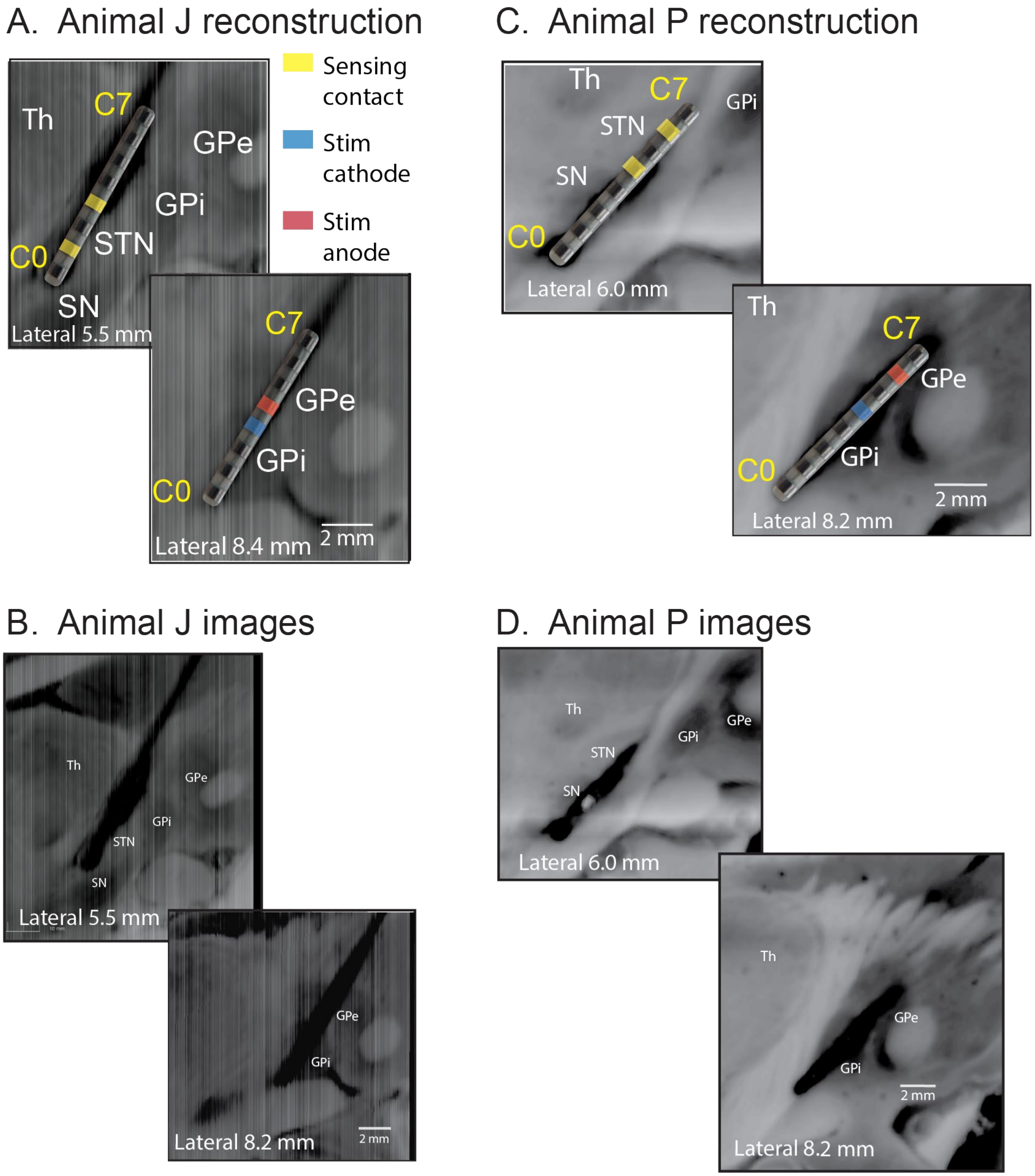


**Fig. A.1.** Localization of DBS electrode arrays used to implement closed-loop stimulation evoked interference (CL-SEI) in animal J and P. Histological reconstructions were created for animal J and P and used to calculate the DBS lead locations in the STN and GPi. The GPi contacts selected for stimulation in the closed-loop system in animal J were C3-4, and in animal P were C4-C6. The STN contacts used for sensing in the closed-loop system were C1-C3 in animal J and C4-C6 in animal P.

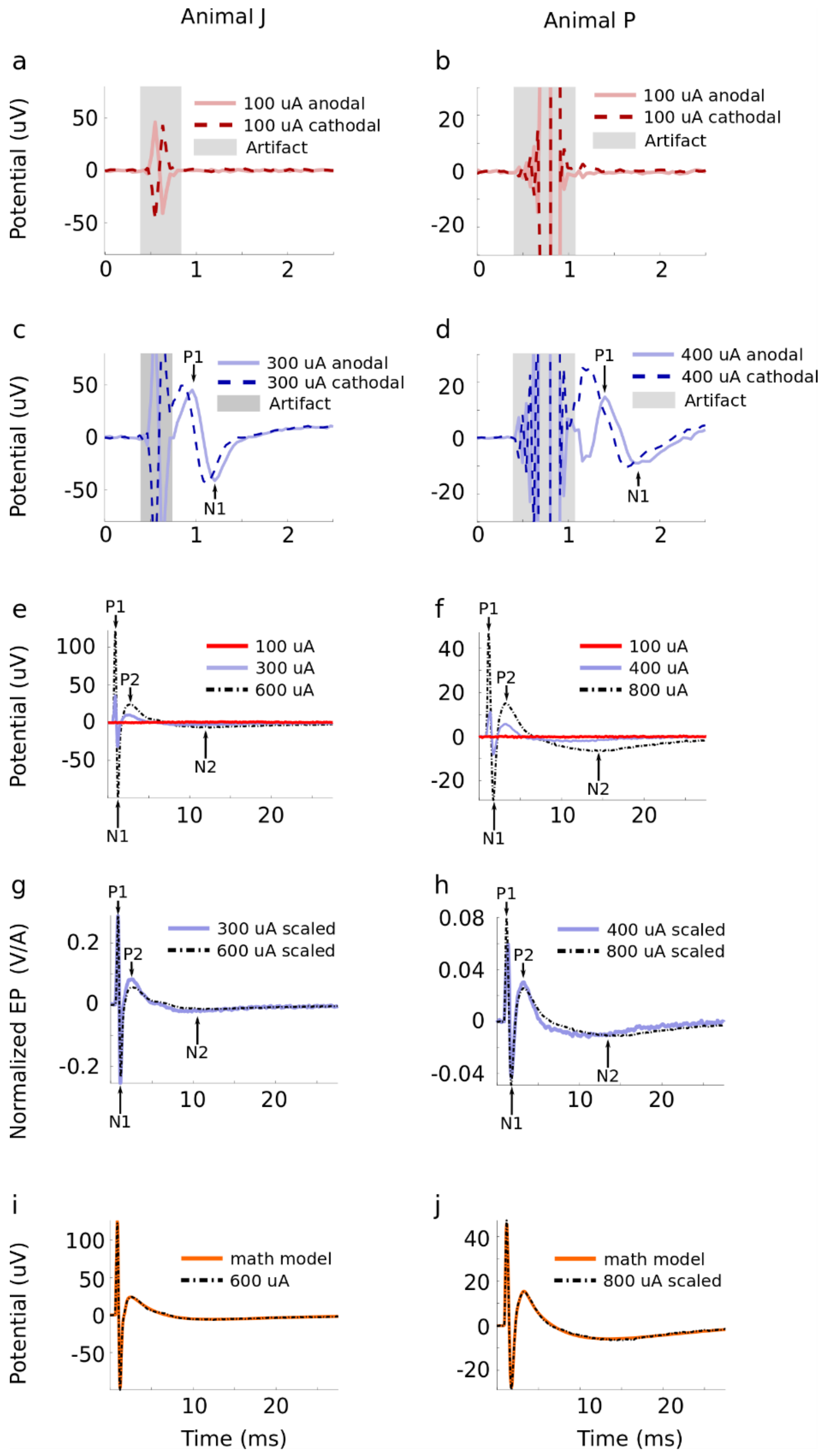


**Fig. A.2.** Characterization and mathematical modeling of potentials in the STN evoked by stimulation in the GPi in animal J and P. (a, b) Sub-threshold potentials for cathodal and anodal stimulation of amplitude 100 μA in both animal J and P. (c, d) Example of super-threshold potentials evoked by cathodal and anodal stimulation with amplitude equal to 300 μA in animal J and 400 μA in animal P. The cathodal response leads the anodal response. (e, f) Evoked potentials generated by stimulation with amplitudes equal to 100, 300, and 600 μA in animal J and 100, 400, and 800 μA in animal P. The peaks (P1, P2) and troughs (N1,

N2) of the two first oscillation cycles are depicted. The evoked potentials are the average response to at least 83 stimulation pulses in animal J and 482 stimulation pulses in animal P for all current amplitudes considered. (g, h) Scaling of evoked potentials by dividing the evoked potentials by the factor $(\boldsymbol{A} - \overline{\boldsymbol{A}})$, where $\boldsymbol{A}$ is the stimulation amplitude and $\overline{\boldsymbol{A}}$ is the minimum current estimated to evoke neural oscillations. The match between the scaled evoked potentials indicate that the scaling approach used in the modeling is accurate to characterize the relationship between the amplitudes of the stimulation pulses and evoked potentials. (i, j) Mathematical models of evoked potentials obtained using system identification techniques shown together with the time series of the evoked potential obtained experimentally in animal J and P.

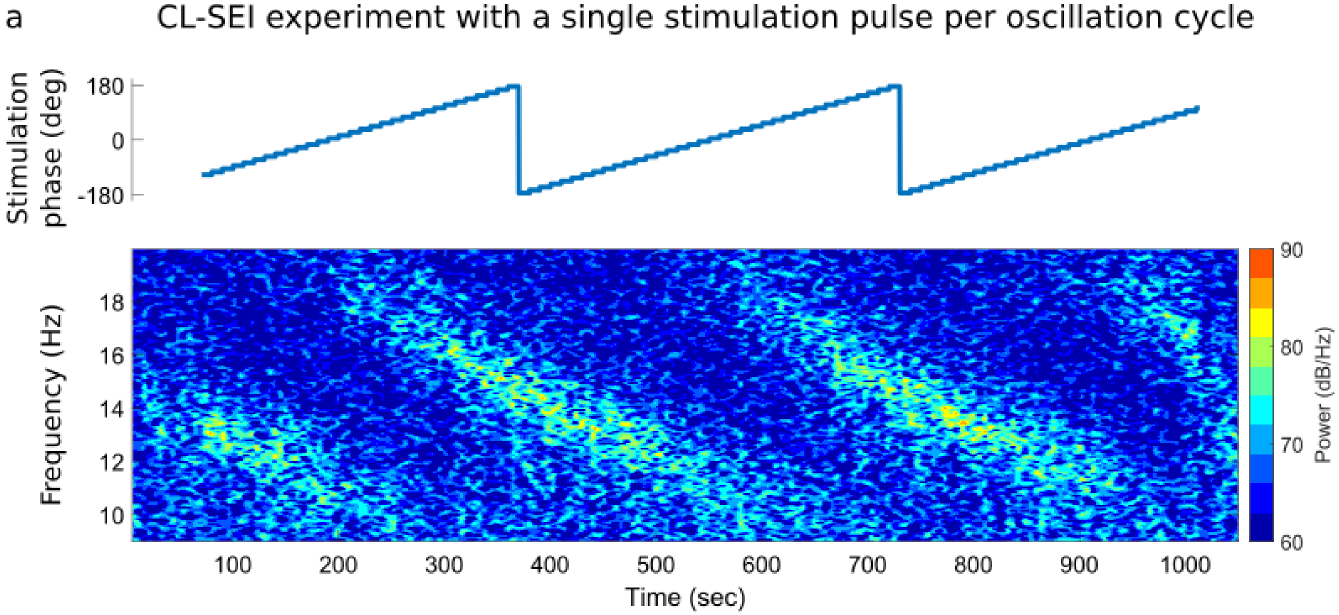


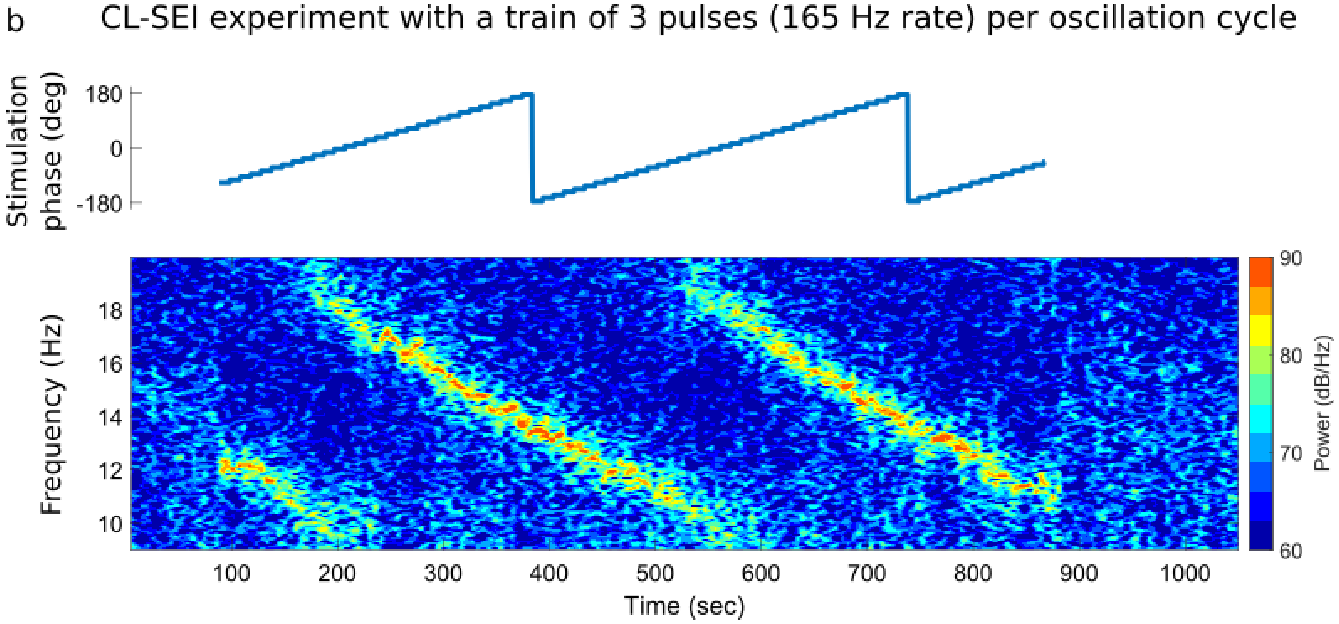

**Fig. A3.** Temporal summation achieved when using a train of three stimulation pulses (b) instead of a single pulse (a) to modulate neural oscillations via CL-SEI. The experimental data was collected from subject P using a stimulation amplitude equal to 600 uA and a pulse width of 80 us. The intra burst frequency of the pulse trains used to generate the data shown in (b) was equal to 165 Hz.

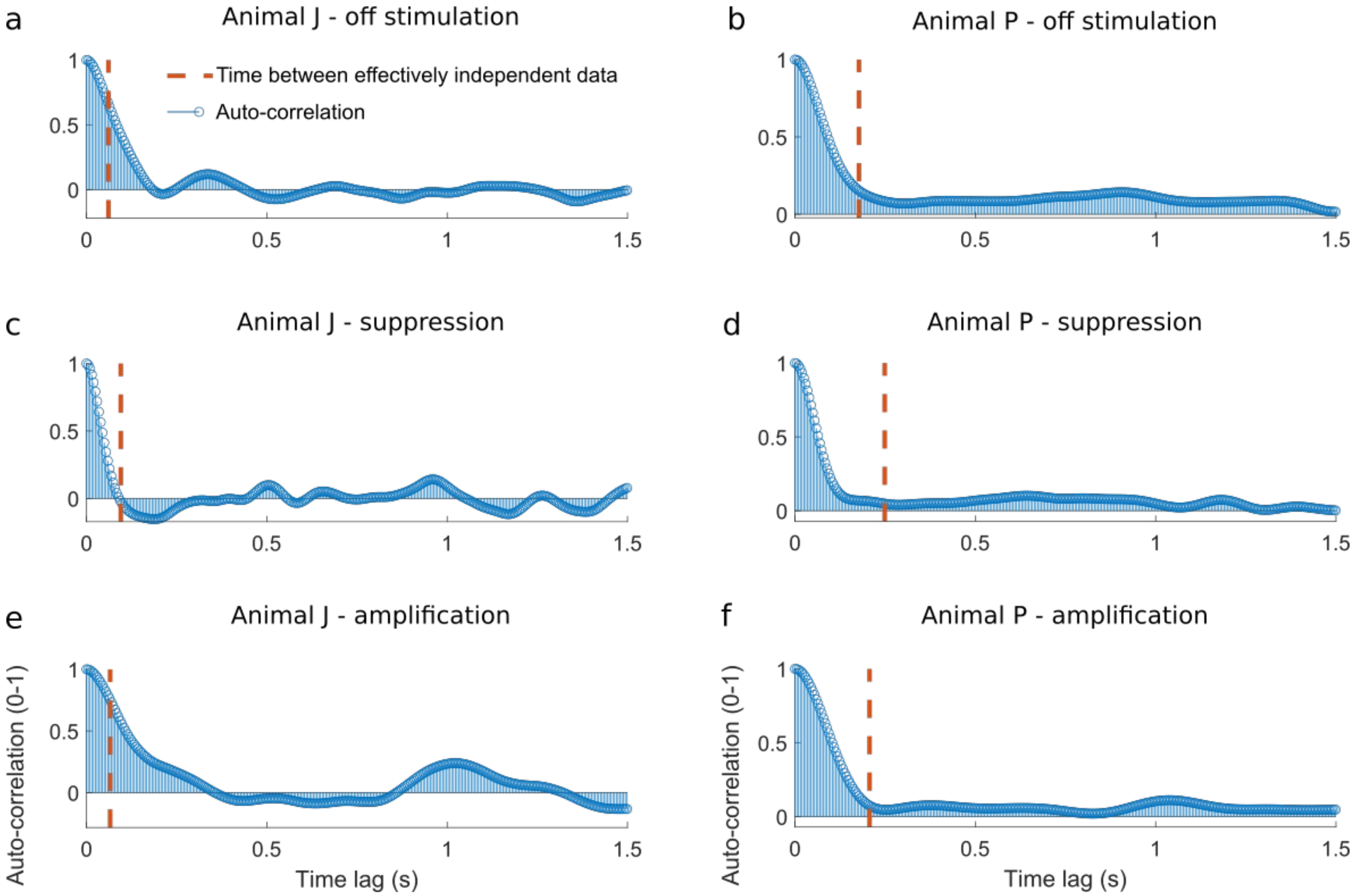


**Fig. A4.** Autocorrelation functions and times between effectively independent data for subject J and P in the off-stimulation, suppression and amplification condition. The maximum time between effectively independent data was 0.25 s (animal P, suppression condition).
